# PrimeDesign software for rapid and simplified design of prime editing guide RNAs

**DOI:** 10.1101/2020.05.04.077750

**Authors:** Jonathan Y. Hsu, Andrew V. Anzalone, Julian Grünewald, Kin Chung Lam, Max W. Shen, David R. Liu, J. Keith Joung, Luca Pinello

**Affiliations:** Department of Biological Engineering, Massachusetts Institute of Technology, Cambridge, MA, USA; Molecular Pathology Unit, Massachusetts General Hospital, Charlestown, MA, USA; Center for Cancer Research and Center for Computational and Integrative Biology, Massachusetts General Hospital, Charlestown, MA, USA; Department of Pathology, Harvard Medical School, Boston, MA, USA; Merkin Institute of Transformative Technologies in Healthcare, Broad Institute of Harvard and MIT, Cambridge, MA, USA; Department of Chemistry and Chemical Biology, Harvard University, Cambridge, MA, USA; Howard Hughes Medical Institute, Harvard University, Cambridge, MA, USA; Computational and Systems Biology Program, Massachusetts Institute of Technology, Cambridge, MA, USA

## Abstract

Prime editing (PE) is a versatile genome editing technology, but design of the required guide RNAs is more complex than for standard CRISPR-based nucleases or base editors. Here we describe PrimeDesign, a user-friendly, end-to-end web application and command-line tool for the design of PE experiments. PrimeDesign can be used for single and combination editing applications, as well as genome-wide and saturation mutagenesis screens. Using PrimeDesign, we also constructed PrimeVar, the first comprehensive and searchable database for prime editing guide RNA (pegRNA) and nicking sgRNA (ngRNA) combinations to install or correct >68,500 pathogenic human genetic variants from the ClinVar database.

## Main text

Prime editing is a new class of mammalian cell genome editing technology that enables unprecedented precision in the installation of specific substitutions, insertions, and deletions into the genome^1^, offering greater versatility than CRISPR nucleases^2,3,4^ and base editors^5,6^. The most efficient prime editing system described to date (referred to as PE3) consists of three components: a fusion protein of a CRISPR-Cas9 nickase and an engineered reverse transcriptase (**RT**), a prime editing guide RNA (**pegRNA**), and a nicking sgRNA (**ngRNA**) (**Supplementary Fig. 1**). The pegRNA targets the Cas9 nickase-RT fusion to a specific genomic locus, but also hybridizes to the nicked single-stranded DNA non-target strand (**NTS**) within the Cas9-induced R-loop, and serves as a template for reverse transcription to create the “flap” that mediates induction of precise genetic changes (**Supplementary Fig. 1a-c**). The ngRNA directs the Cas9 nickase-RT fusion to nick the target strand (i.e. the strand opposite the flap) and thereby biases repair towards the desired change encoded in the flap (**Supplementary Fig. 1d-e**).

The complexity of the PE3 system makes it time-consuming to manually design the required pegRNA and ngRNA components. Beyond the need to design the spacer for both guide RNAs, there are multiple other parameters that must be accounted for that can impact their editing efficiencies, including: primer binding site (**PBS**) length, reverse transcription template (**RTT**) length, and distance between the pegRNA and ngRNA. Here we present PrimeDesign, a user-friendly web application (http://primedesign.pinellolab.org/) (**Fig. 1**) and command-line tool (https://github.com/pinellolab/PrimeDesign) that automates and thereby simplifies the design of pegRNAs and ngRNAs for single edits, combination edits, and genome-wide and saturation mutagenesis screens.

**Figure 1:**
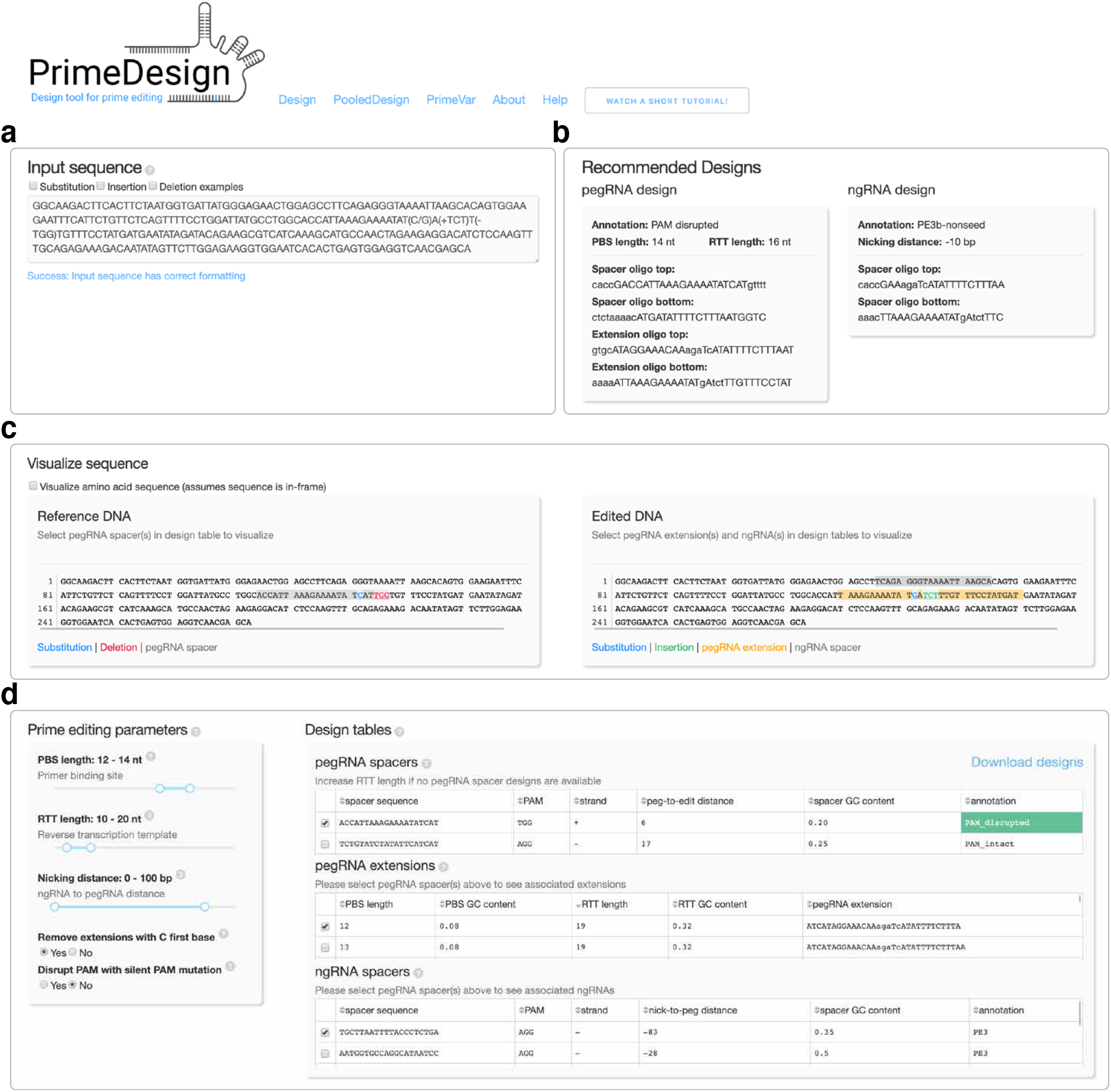
PrimeDesign web application. **a)** PrimeDesign takes a single sequence as input encoding both the original reference and desired edited sequences, **b)** recommends a candidate pegRNA and ngRNA combination to install the edit of interest, **c)** provides sequence visualizations of the edit of interest and selected pegRNA and ngRNA designs, and **d)** enables the interactive design of both pegRNAs and ngRNAs that can be downloaded as a summary table.

PrimeDesign uses a single input that encodes both the original reference and the desired edited sequences (**Fig. 1a**, **Supplementary Note 1**), recommends a candidate pegRNA and ngRNA combination to install the edit of interest (**Fig. 1b**), provides sequence visualization of the prime editing event (**Fig. 1c**), and enumerates all possible pegRNA spacers, pegRNA extensions, and ngRNAs within optimized parameter ranges (as previously defined by the Liu group^1^) for installing the desired edit (**Fig. 1d**). PrimeDesign provides important annotations for pegRNA (e.g. PAM disrupted) and ngRNA (e.g. PE3b) designs and streamlines the design of PAM-disrupting silent mutations to improve editing efficiency and product purity (**Supplementary Note 2**). In addition, PrimeDesign enables the pooled design of pegRNA and ngRNA combinations for genome-wide and saturation mutagenesis screens (http://primedesign.pinellolab.org/pooled), and ranks the designs according to best design practices^1^. The saturation mutagenesis feature allows for mutagenesis at single-base or single-amino acid resolution; PrimeDesign automatically constructs all edits within a user-defined sequence range and generates the designs to install these edits (**Supplementary Note 3**).

To demonstrate the utility of PrimeDesign, we took pathogenic human genetic variants from ClinVar^7^ (n= 69,481) and designed candidate pegRNAs and ngRNAs for the correction of these pathogenic alleles. Of these pathogenic variants, we found that 91.7% are targetable by at least a single pegRNA spacer with a maximum RTT length of 34 nt (**Fig. 2a** and **Supplementary Table 1**). An average of 3.7 pegRNA spacers were designed per pathogenic variant, representing multiple options for prime editing to correct each variant. Furthermore, 25.9% of targetable pathogenic variants included at least a single pegRNA that disrupts the PAM sequence, which has been associated with improved editing efficiency and product purity. The PE3b strategy (the design of ngRNAs that preferentially nick the non-edited strand *after* edited strand flap resolution) is viable for 79.5% of targetable variants (59.7% when only considering mismatches in the seed sequence) (**Fig. 2b** and **Supplementary Table 1**). Lastly, 11.9% of targetable pathogenic variants are amenable to both the PAM-disrupting and PE3b seed-mismatched strategies (**Supplementary Table 1**).

**Figure 2:**
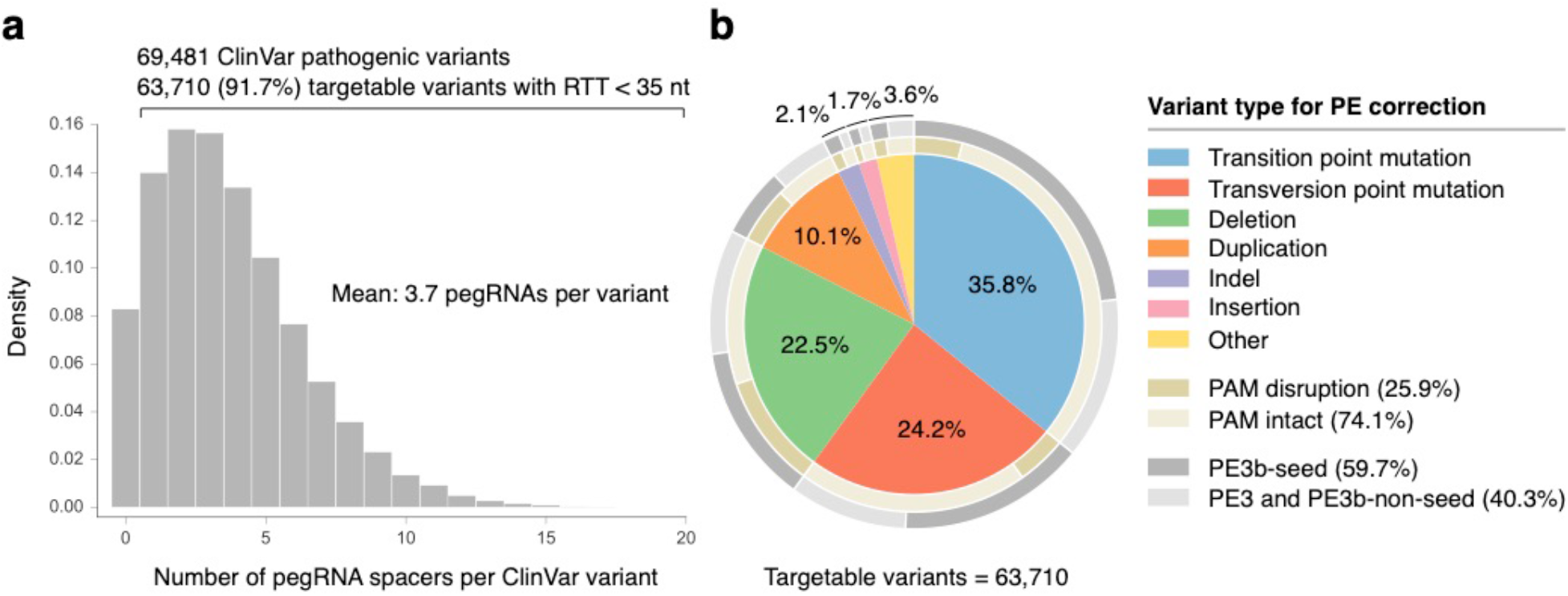
PrimeDesign analysis of the ClinVar database. **a)** The distribution of the number of designed pegRNA spacers per ClinVar variant. Candidate pegRNAs were determined based on the requirement of RTT length <35 nt and the RT extension to have a minimum homology of 5 nt downstream of the edit. **b)** The 63,710 (91.7%) targetable ClinVar variants classified by type. The inner ring (gold) represents the proportion of targetable variants by type where at least one pegRNA could be designed to disrupt the PAM sequence (dark gold). The outer ring (grey) represents the proportion of targetable variants by type where at least one ngRNA could be designed for the PE3b strategy where the mismatch lies in the seed sequence (PAM-proximal nucleotides 1-10) (dark grey). See Supplementary Table 1 for details.

To make all of the ClinVar prime editing designs more accessible, we constructed PrimeVar (http://primedesign.pinellolab.org/primevar), a comprehensive and searchable database for pegRNA and ngRNA combinations to install or correct >68,500 pathogenic human genetic variants. Using either the dbSNP reference SNP number (rs#) or ClinVar Variation ID, candidate pegRNAs and ngRNAs are readily-available across a range of PBS (10-17 nt) and RTT (10-80 nt) lengths.

In summary, PrimeDesign is a comprehensive and general method for facile and automated design of pegRNAs and ngRNAs. We note that optimal pegRNAs are target site-dependent^1^, and thus researchers may still need to refine pegRNA choices after testing designs proposed by PrimeDesign. Nonetheless, we envision that PrimeDesign should greatly simplify the design of prime editing components and thereby increase the use and accessibility of this powerful and important technology^8,9^.

---

**Supplementary Figure 1:**
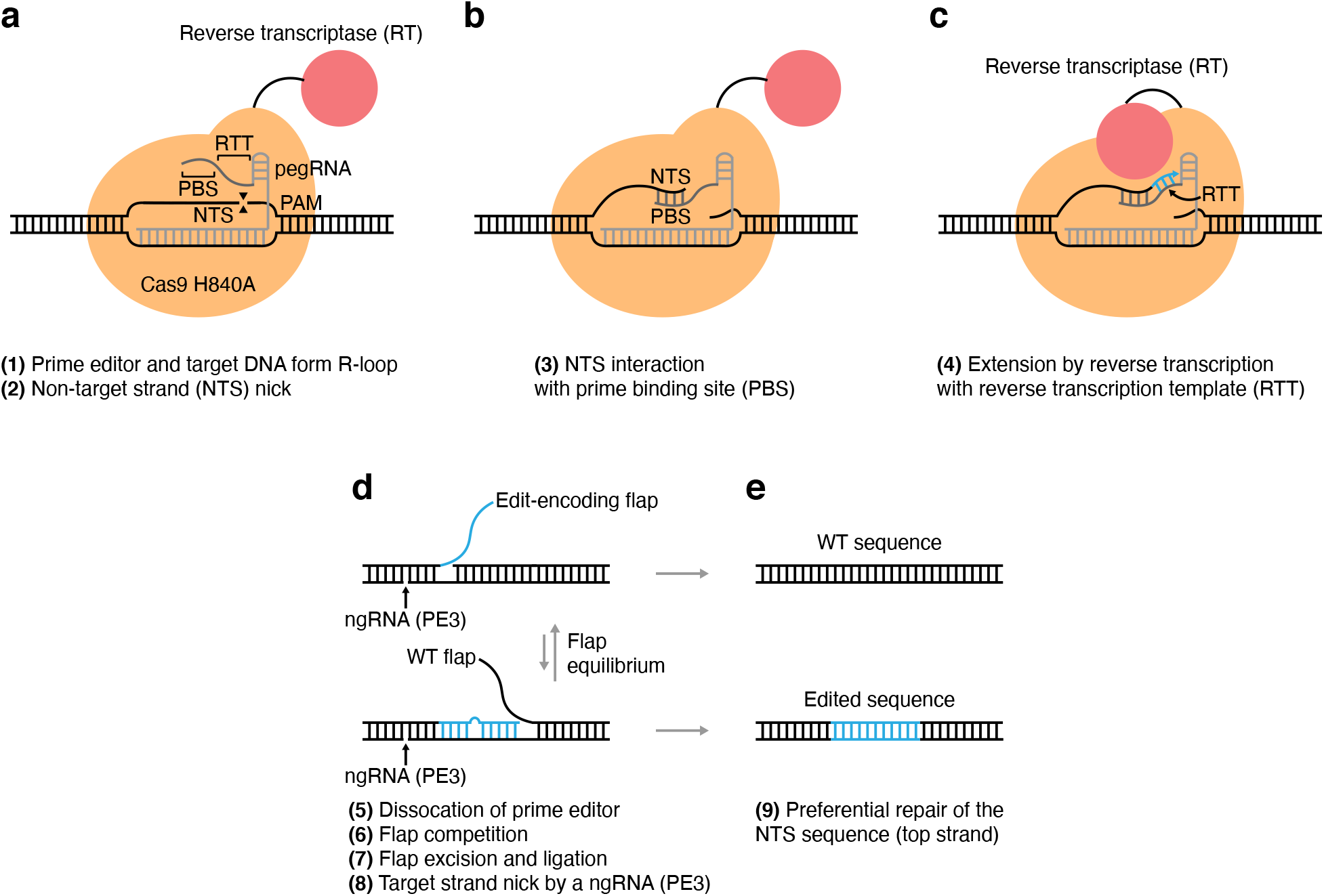
Overview of prime editing. **a)** The prime editor, a fusion protein of a CRISPR-SpCas9 H840A nickase and an engineered Moloney murine leukemia virus reverse transcriptase (MMLV-RT), coupled with a prime editing guide RNA (pegRNA) engages target DNA to form an R-loop and introduces a nick in the non-target strand (NTS). **b)** The 3’ end of the NTS interacts with the 3’ end of the pegRNA extension, called the primer binding site (PBS), and this stabilized interaction allows for **c)** extension of the NTS by reverse transcription with the reverse transcriptase and reverse transcription template (RTT) of the pegRNA. **d)** The prime editor system dissociates from its target DNA and the newly-synthesized edit-encoding strand competes with the wild-type (WT) strand for hybridization; flap excision and ligation occurs, and a target strand (TS) nick is introduced with a nicking sgRNA (ngRNA) for the PE3 strategy. **e)** The NTS sequence is preferentially repaired with the TS nick, resulting in a bias towards successful editing in scenarios where the edit-encoding flap outcompetes the WT flap for genomic incorporation.

## Supplementary Note 1

The PrimeDesign input sequence format encodes both the original reference and desired edited sequences. All edits are formatted within a set of parentheses. Substitution edits are encoded by: (*ref*/*edit*), where *ref* is the pre-substitution reference sequence and *edit* is the post-substitution edited sequence. Insertion edits are encoded by: (+*ins*) or (/*ins*), where *ins* is the sequence to be inserted into the reference sequence during the editing event. Deletion edits are encoded by: (-*del*) or (*del*/), where *del* is the sequence to be deleted from the reference sequence during the editing event. Substitution, insertion, and deletion edits can be combined for combination edits. All sequences unaffected by editing remain outside of the parentheses. It is recommended to place the intended edit site near the center of the input sequence and have the total input sequence length be >300 bp to ensure thorough design for prime editing. We provide some examples of input sequences below:

*Substitution edit:*

CACACCTACACTGCTCGAAGTAAATATGCGAAGCGCGCGGCCTGGCCGGAGGCGTTCCGCGCCGCCACGTGTTCGTTAACTGTTGATTGGTGGCACATAAGCAATCGTAGTCCGTCAAATTCAGCTCTGTTATCCCGGGCGTTATGTGTCAAATGGCGTAGAACGGGATTGACTGTTTGACGGTAGCTGCTGAGGCGG(G/T)AGAG ACCCTCCGTCGGGCTATGTCACTAATACTTTCCAAACGCCCCGTACCGATGCTGAACAAGTCGATGCAGGCTCCCGTCTTTGAAAAGGGGTAAACATACAAGTGGATAGATGATGGGTAGGGGCCTCCAATACATCCAACACTCTACGCCCTCTCCAAGAGCTAGAAGGGCACCCTGCAGTTGGAAAGGG

*Insertion edit:*

CACACCTACACTGCTCGAAGTAAATATGCGAAGCGCGCGGCCTGGCCGGAGGCGTTCCGCGCCGCCACGTGTTCGTTAACTGTTGATTGGTGGCACATAAGCAATCGTAGTCCGTCAAATTCAGCTCTGTTATCCCGG GCGTTATGTGTCAAATGGCGTAGAACGGGATTGACTGTTTGACGGTAGCTGCTGAGGCGGGA(+GTAA) GAGACCCTCCGTCGGGCTATGTCACTAATACTTTCCAAACGCCCCGTACCGATGCTGAACAAGTCGATGCAGGCTCCCGTCTTTGAAAAGGGGTAAACATACAAGTGGATAGATGATGGGTAGGGGCCTCCAATACATCCAACACTCTACGCCCTCTCCAAGAGCTAGAAGGGCACCCTGCAGTTGGAAAGGG

*Deletion edit:*

CACACCTACACTGCTCGAAGTAAATATGCGAAGCGCGCGGCCTGGCCGGAGGCGTTCCGCGCCGCCACGTGTTCGTTAACTGTTGATTGGTGGCACATAAGCAATCGTAGTCCGTCAAATTCAGCTCTGTTATCCCGGGCGTTATGTGTCAAATGGCGTAGAACGGGATTGACTGTTTGACGGTAGCTGCTGAGGCGGGAG(-AGAC)CCTCCGTCGGGCTATGTCACTAATACTTTCCAAACGCCCCGTACCGATGCTGAACAAGTCGATGCAGGCTCCCGTCTTTGAAAAGGGGTAAACATACAAGTGGATAGATGATGGGTAGGGGCCTCCAATACATCCAACACTCTACGCCCTCTCCAAGAGCTAGAAGGGCACCCTGCAGTTGGAAAGGG

*Combination edit:*

CACACCTACACTGCTCGAAGTAAATATGCGAAGCGCGCGGCCTGGCCGGAGGCGTTCCGCGCCGCCACGTGTTCGTTAACTGTTGATTGGTGGCACATAAGCAATCGTAGTCCGTCAAATTCAGCTCTGTTATCCCGGGCGTTATGTGTCAAATGGCGTAGAACGGGATTGACTGTTTGACGGTAGCTGCTGAGGCGG(G/T)A(+GT AA)G(-AGAC)CCTCCGTCGGGCTATGTCACTAATACTTTCCAAACGCCCCGTACCGATGCTGAACAAGTCGATGCAGGCTCCCGTCTTTGAAAAGGGGTAAACATACAAGTGGATAGATGATGGGTAGGGGCCTCCAATACATCCAACACTCTACGCCCTCTCCAAGAGCTAGAAGGGCACCCTGCAGTTGGAAAGGG

## Supplementary Note 2

PrimeDesign provides important annotations during the design of pegRNAs and ngRNAs. For pegRNAs, the different annotations include: *PAM intact*, *PAM disrupted*, and *PAM disrupted silent mutation*. The *PAM intact* annotation is given to pegRNAs that do not introduce edits into the PAM sequence at positions that have sequence preference, whereas the *PAM disrupted* annotation is given to pegRNAs that introduce sequence modifications at PAM positions that have sequence preference. For coding sequence edits, PrimeDesign offers a functionality to introduce silent mutations to potentially improve editing efficiency and product purity. When this functionality is turned on and the design is available, the *PAM disrupted silent mutation* is provided for suitable pegRNA designs. Importantly, the input sequence must be provided in-frame in order for this function to work properly. We recommend using the amino acid sequence viewer on our PrimeDesign web application to check whether the input sequence is in-frame, and deleting the left-most bases of the input sequence to achieve the correct frame. PrimeDesign uses the GenScript human codon usage frequency table (https://www.genscript.com/tools/codon-frequency-table) and automatically selects the best codon by frequency to introduce the silent mutation.

For ngRNAs, the different annotations include: *PE3*, *PE3b non-seed*, and *PE3b seed*. The *PE3* annotation is given to ngRNAs that have a spacer match to both the original reference and desired edited sequences. The *PE3b non-seed* and *PE3b seed* annotations are given to ngRNAs that have a spacer that only perfectly matches the desired edited sequence, and therefore preferentially nick the non-edited strand *after* edited strand flap resolution. The *PE3b non-seed* ngRNAs contain sequence mismatches to the original reference sequence outside of PAM-proximal nucleotides 1-10 (seed region), whereas *PE3b seed* ngRNAs contain sequence mismatches to the original reference sequence within the seed region. Spacer mismatches in the seed region severely inhibit target DNA binding and cleavage, and do so to a larger degree compared to spacer mismatches outside of the seed region. For this reason, *PE3b seed* ngRNAs may exhibit higher specificity in nicking the non-edited strand *after* edited strand flap resolution and are therefore more suitable for the PE3b strategy compared to *PE3b non-seed* ngRNAs.

## Supplementary Note 3

PrimeDesign offers the ability to perform pooled designs for genome-wide and saturating mutagenesis screen applications. For each edit of interest, PrimeDesign outputs a user-defined number of pegRNAs (unique spacers) and ngRNAs per pegRNA. These designs are ranked according to general guidelines previously established by the Liu group. Hierarchal ranking of pegRNAs is performed by first using the pegRNA annotations (*PAM disrupted* -> *PAM disrupted silent mutation* -> *PAM intact*), and then using pegRNA-to-edit distances (smallest to largest). Hierarchal ranking of ngRNAs is performed by first using the ngRNA annotations (*PE3b seed* -> *PE3b non-seed* -> *PE3*), and then using deviations from a user-defined ngRNA-to-pegRNA distance parameter (default: 75 bp).

PrimeDesign enables streamlined design of saturation mutagenesis studies with prime editing at single-base and single-amino acid resolution. The PrimeDesign input sequence format for saturation mutagenesis applications is the following: (*seq*), where *seq* is the user-defined sequence range of where the saturating mutagenesis will take place. If the “base” option is selected, PrimeDesign will automatically construct all single-base changes (i.e. A -> T,C,G) across the user-defined sequence range. If the “amino acid” option is selected, PrimeDesign will automatically construct all single-amino acid changes (including a stop codon) within the user-defined sequence range. Importantly, the user-defined sequence range must be in-frame in order for this function to work properly. PrimeDesign uses the GenScript human codon usage frequency table (https://www.genscript.com/tools/codon-frequency-table) and automatically selects the best codon by frequency to introduce the amino acid changes.

## Code Availability Statement

The PrimeDesign command-line tool and web application code can be found at https://github.com/pinellolab/PrimeDesign.

## Acknowledgements

L.P. is supported by the National Human Genome Research Institute (NHGRI) Career Development Award (R00HG008399), Genomic Innovator Award (R35HG010717) and CEGS RM1HG009490. J.K.J. is supported by NIH R35 GM118158, NIH RM1 HG009490, the Robert B. Colvin, M.D. Endowed Chair in Pathology, and the Desmond and Ann Heathwood MGH Research Scholar Award. D.R.L. is supported by the Merkin Institute of Transformative Technologies in Healthcare, US NIH grants U01AI142756, RM1HG009490, R01EB022376, and R35GM118062, and the HHMI. A.V.A. acknowledges a Jane Coffin Childs postdoctoral fellowship. J.G. was funded by the Deutsche Forschungsgemeinschaft (DFG, German Research Foundation) – Projektnummer 416375182.

## Author Contributions

J.Y.H. developed PrimeDesign. A.V.A., J.G., and K.C.L provided feedback during the development of PrimeDesign. M.W.S. contributed to the ClinVar analysis. L.P., J.K.J., and D.R.L. supervised the project and provided feedback and guidance. J.Y.H., L.P., J.K.J., and D.R.L. wrote the manuscript with input from all other authors.

## Methods

### PrimeDesign analysis on ClinVar variants

The ClinVar database was accessed April 8^th^ 2020. Variants were filtered with the following conditions: 1) included a valid GRCh38/hg38 coordinate 2) labeled as *Pathogenic* for the column “ClinicalSignificance” 3) contained a unique identifier determined by the concatenation of columns “Name,” “RS# (dbSNP),” and “VariationID.” All variants with ambiguous IUPAC code were converted into separate entries with non-ambiguous bases for downstream analysis. Following these steps, the total number of ClinVar variants totaled 69,481. Sequence inputs were formatted for all entries for both the installation and correction of these pathogenic variants. After running PrimeDesign on the ClinVar variants, candidate pegRNA designs were filtered with two criteria: 1) maximum RTT length of 34 nt and 2) minimum homology of 5 nt downstream of the edit. The pegRNAs with *PAM disrupted* annotations have mutations in the dinucleotide GG of the NGG motif, and the ngRNAs with *PE3b*, *PE3b non-seed*, and *PE3b seed* annotations have mismatches anywhere in the protospacer, mismatches outside of PAM-proximal nucleotides 1-10, or mismatches within PAM-proximal nucleotides 1-10, respectively.

### Construction of PrimeVar database

The filtered ClinVar variants from the PrimeDesign analysis were used to build a comprehensive database of candidate pegRNA and ngRNA combinations. Prime editing designs are available to install and correct the pathogenic human genetic variants. PrimeDesign was run with a PBS length range of 10-17 nt, RTT length range of 10-80 nt, and ngRNA distance range of 0-100 bp. All of the pegRNA and ngRNA designs for each variant are stored on PrimeVar (http://primedesign.pinellolab.org/primevar).

